# Acute Triclosan Exposure Alters Mitochondrial Form and Function in *Xenopus laevis* Tadpoles, Including the Developing Brain

**DOI:** 10.1101/2024.11.01.621554

**Authors:** Alex H Thompson, Christopher K Thompson

## Abstract

Mitochondrial health is critical for normal brain development, and mitochondrial toxicants impair energy production and induce mitophagy, resulting in network fission. This can substantially impact cellular and molecular pathways essential for brain development. Recent studies have shown that triclosan, a once widely used antibacterial and antifungal agent, acts as a mitochondrial uncoupler. However, the impact of triclosan on mitochondrial function in the developing brain has yet to be thoroughly investigated. We performed a series of experiments designed to assess the effects of triclosan on mitochondrial form and function in the developing *Xenopus laevis* tadpole brain. We injected tadpole brains with TMRM, a fluorescent reporter of mitochondrial membrane potential. After two hours, we imaged the brain using high-resolution confocal microscopy to obtain a baseline level of TMRM fluorescence, then immediately treated tadpoles with either triclosan (10uM, 5uM, 1uM, 0.5uM 0.1uM), FCCP (0.5uM), or control, and then continued imaging the brain once every five min for 30 min. We found that 10uM triclosan significantly decreased TMRM fluorescence in mitochondria in the end feet of radial glial cells in ∼ 15-20 min; lower concentrations operated over a longer time course. We also observed that mitochondrial networking was disrupted as they underwent fission. We also measured metabolic function in vivo in whole tadpoles using an XF Seahorse Flux Analyzer, acutely exposing tadpoles to either triclosan (30uM, 10uM, 1uM) or control. Triclosan immediately increased oxygen consumption rates dose-dependently, suggesting mitochondrial uncoupling. These results show that triclosan has the capacity to impair mitochondrial form and function in developing neural tissue.

## Introduction

Triclosan (TCS) was first introduced in 1970 as an antimicrobial additive; where it eventually entered thousands of health and home care products, including deodorants, mouthwash, toothpaste, cosmetics, shampoo, skin creams, plastic kitchenware, children’s products, shoes, and textiles. Companies marketed TCS with claims of prolonged shelf life and increased cleanliness [1]. Yet TCS has been implicated in many adverse health outcomes, including impaired metabolism and even some aspects of cognitive function [2]. After an FDA risk assessment, TCS was banned from commercial soaps [3]. It is still widely prevalent in other goods and products, with no explicit medical claim on the product’s label [4]. Before the FDA decision, TCS was among the top ten most commonly detected organic wastewater compounds in the United States [5–8] and is readily absorbed through the skin [9]. TCS is still seen in human urine, breastmilk, blood, and newborn’s umbilical cord at astonishing rates [10–16]. Children born to mothers who had detectable levels of TCS in their urine at the child’s birth have been associated with a decrease in IQ score, verbal comprehension, and perceptual reasoning when tested at eight years-of-age [17]. This implies that TCS may impair brain development, and a study in rats showed that there were autism-like, repetitive symptoms in the adult offspring exposed to TCS during pregnancy [18]. Some have suggested that the increased exposure to TCS in the last 20+ years may explain the growing number of autism spectrum disorder (ASD) cases. Autism rates rose from 1/149 children in 2000 to about 1/40 children in 2016 [19], with the mean age of first birth only changing by roughly 2.5 years in that same time [20]. Epidemiological data suggest that an environmental component likely explains, at least in part, some of the increase in incidence.

Only a handful of studies have examined the effects of TCS exposure on the brain. In the snail nervous system, TCS significantly decreases action potential amplitude. Where in mouse neuromuscular end junctions, TCS increases spontaneous vesicle release and reduces membrane potential [21]. Additionally, decreased neural synaptic density and increased apoptosis are seen in the brains of developing zebrafish after a TCS treatment [22]. To fill in some of the gaps in our understanding of how TCS affects the brain, we conducted several experiments to examine the effects of TCS on mitochondrial function in *Xenopus laevis* tadpoles, including investigations to determine how TCS affects neural progenitor cells (NPCs, also known as radial glial cells), see Figure 1 for reference. These cells divide and give rise to neurons, and they may be especially susceptible to TCS because neuronal proliferation is controlled by mitochondrial activity within the cell [23]. A growing body of evidence indicates a long-term, detrimental effect on the mitochondria due to TCS’s lipophilic properties directly targeting the phospholipid membrane. TCS disrupts mitochondrial form and function through its uncoupling properties [24], which we hypothesize as its primary influence on the disruption of the proton gradient. At this point, few studies explore TCS’s effects on mitochondrial changes to NPCs and overall physiological changes *in vivo*.

**Figure 1:**
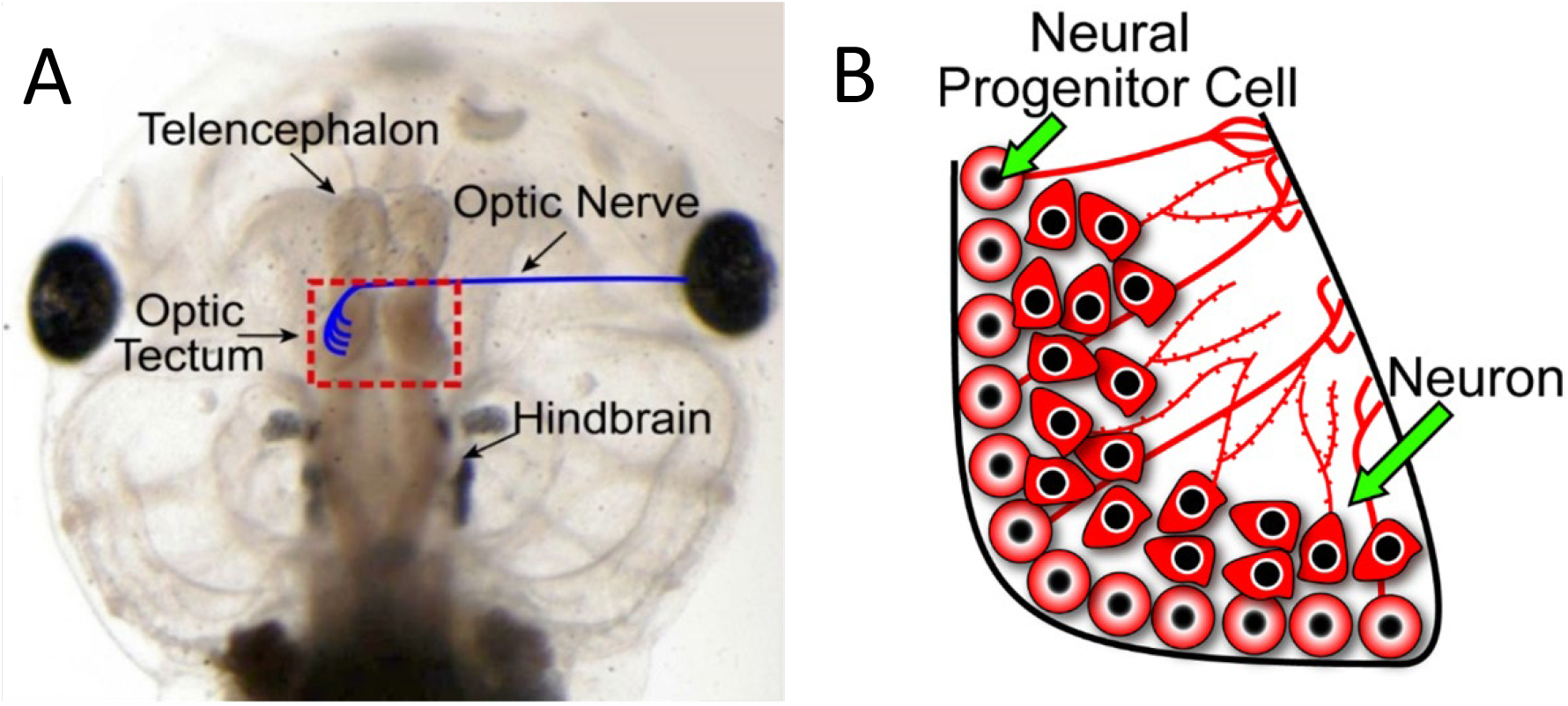
A) Dorsal aspect of the head of a Xenopus laevis tadpole. B) A schematic diagram of the right hemisphere of the midbrain area in the red box in A illustrating the retinotectal circuit. Neural progenitor cells are found in the ventricular proliferative zone. Neurons are generated from NPCs. They extend dendrites into the neuropil, forming synapses with other tectal neurons and/or RGC axons (blue in panel A).

To further investigate the toxicological effects of TCS in the developing brain *in vivo*, we used *Xenopus laevis* tadpoles (also known as the African clawed frog). Tadpoles are an ideal animal model because we can tightly control environmental concentrations of TCS, as well as monitor morphological changes as the brain rapidly develops. Additionally, due to the tadpole’s transparent skin over the brain, we can observe the real-time response to TCS in individual cells and even in specific mitochondria. This model was suitable to test the hypothesis of TCS impairing mitochondrial function in developing tissues. Additional investigations into the mechanism of action of TCS are essential to uncovering ways of ensuring no further damage to future populations and aiding in possible approaches to reversing the impairments.

## Methods

### Chemicals and Reagents

TCS (Irgasan, CAS 3380-34-5, 97.0%),

FCCP (Carbonyl cyanide 4-(trifluoromethoxy)phenylhydrazone, CAS 370-86-5, 98%)

Oligomycin

Antimycin A

MS222 (Tricaine methanesulfonate, CAS 886-86-2, 98%)

was purchased from Sigma-Aldrich (St. Louis, Mo, USA).

-60x E3 Media: 297.7413mM NaCL, 10.7309mM KCL, 19.7265mM CaCl2H4O2, 24.0531mM

Cl2H12MgO6, added to Millipore H2O. Using HCL, pH to 7.4.

-0.01% MS-222, 1x E3 Media: 50mL 0.2% MS-222, 16.5 mL of 60× stock E3 medium, brought to 1 liter using Millipore H2O.

### Animal husbandry

*Xenopus laevis* were maintained through established methods [25]. The adult, *Xenopus laevis*, were raised at a temperature of 18°C and were scheduled for controlled feedings. Breeding was induced between two healthy adult frogs with subcutaneous injections of human chorionic gonadotropin. This would yield a clutch of roughly 2000 fertilized eggs overnight. Adult frogs were removed, and embryos were raised to either stage 22-25 (3-4 days old) or tadpoles to stage 46-47 (7-10 days old).

### Mortality Analysis

Treatment groups were established to assess Triclosan’s effects on the overall survivability and range of concentration on the Xenopus laevis tadpoles. Embryos in stage 25 were placed in treatment groups of either vehicle control, 0.5uM FCCP, or a TCS concentration of 0.1uM, 0.5uM, 1uM, 5uM, 10uM, or 50uM. A population of 12 embryos a bowl, at two bowls a treatment, repeated in a separate clutch (n=48). All treatments were diluted in Steinberg’s solution. Survival rates were measured over 14 days and classified based on active heartbeat under a brightfield microscope.

### Mitochondrial labeling

Embryos were incubated in TMRM diluted in Steinburg’s for 2 hours prior to experimentation Mitochondria in the brain were labeled with TMRM via intraventricular injection through a glass micropipette 24 hours before imaging.

### Triclosan Treatment for Imagining

Embryos or tadpoles were placed in TCS (30uM, 10uM, 5uM, 1uM) or positive control FCCP (0.5uM) made in Steinberg’s solution. All experiments shown are acute exposures to TCS. Images were taken every 5 minutes following acute TCS treatment. Images were normalized based on pre-treatment fluorescence intensity.

### Confocal imaging and analysis

Embryos and tadpoles were imaged using a Lecia SP8 confocal microscope; images were analyzed using ImageJ.

### Extracellular Metabolic Flux Analysis

Oxygen consumption rate (OCR) was measured using an XFe24 Seahorse Extracellular Flux Analyzer (Agilent Technologies, Santa Clara, California, USA). The day before planned assay, the XFe24 Extracellular Flux Assay Kit cartage was hydrated using 1mL of XF Calibrant for each 24 wells, then placed into a 25ºC incubator. Healthy Xenopus Laevis eggs were incubated at 25ºC following fertilization. Embryos were incubated till stage 23, roughly 48 hours post-fertilization. Using the hydrated cartridge, either vehicle control, 300uM, 100uM, or 10uM TCS was loaded into well A at a volume of 70uL for the final concentrations of 30uM 10uM or 1uM per well. An XF Cell Mitochondrial Stress Test was prepared to assess the bioenergetic status of the embryos by injecting ATPase inhibitor oligomycin (10uM) in well B, inner membrane uncoupler FCCP (2.5uM) in well C. Complex III inhibitor, antimycin A (1uM) with complex I inhibitor rotenone (1uM) in well D. 22 embryos were loaded into Agilent Seahorse Islet Capture Microplates, with provided micro screens encasing each embryo. Embryos were washed twice with 0.01% MS-222, 1x E3 media before adding the final volume of 630uL to account for increasing volume changes from each injection. An XF Cell Mitochondrial Stress Test was carried out with the heating element turned off on the XFe24 Seahorse Extracellular Flux Analyzer before assay. Data are expressed as percent control of pmol of O_2_ per minute per each well’s stabilized baseline measurement.

### Statistical analysis

Two-way ANOVAs and subsequent graphs were constructed using Graph Pad Prism. These were used to analyze the impact of the two factors (time and dose) on the response variables (e.g., change in TMRM intensity).

## Results

Experimental groups were established to assess the efficacy of Triclosan (TCS) and dosage levels on Xenopus laevis tadpoles. The survival rates of the tadpoles were monitored at regular intervals over 14 days after exposure to varying concentrations of TCS and the positive control, 0.5 um FCCP, to improve our understanding. The results, depicted in Figure 2, indicated that concentrations exceeding 5um TCS were acutely toxic, leading to fatalities within hours for all tested tadpoles. Although FCCP was also found to be lethal, it exhibited delayed toxicity compared with high doses of TCS, which was valuable information for future experiments. While tolerable concentration levels could extend as low as 1 uM for prolonged durations, mortality rates surged drastically during the last day using this dose range. This suggests that caution should be taken when considering use cases with such levels or durations beyond seven days. Interestingly, no significant differences in survivability were observed between control groups and those exposed dosages ranging from 0.1 uM to 0. 5uM.

**Figure 2:**
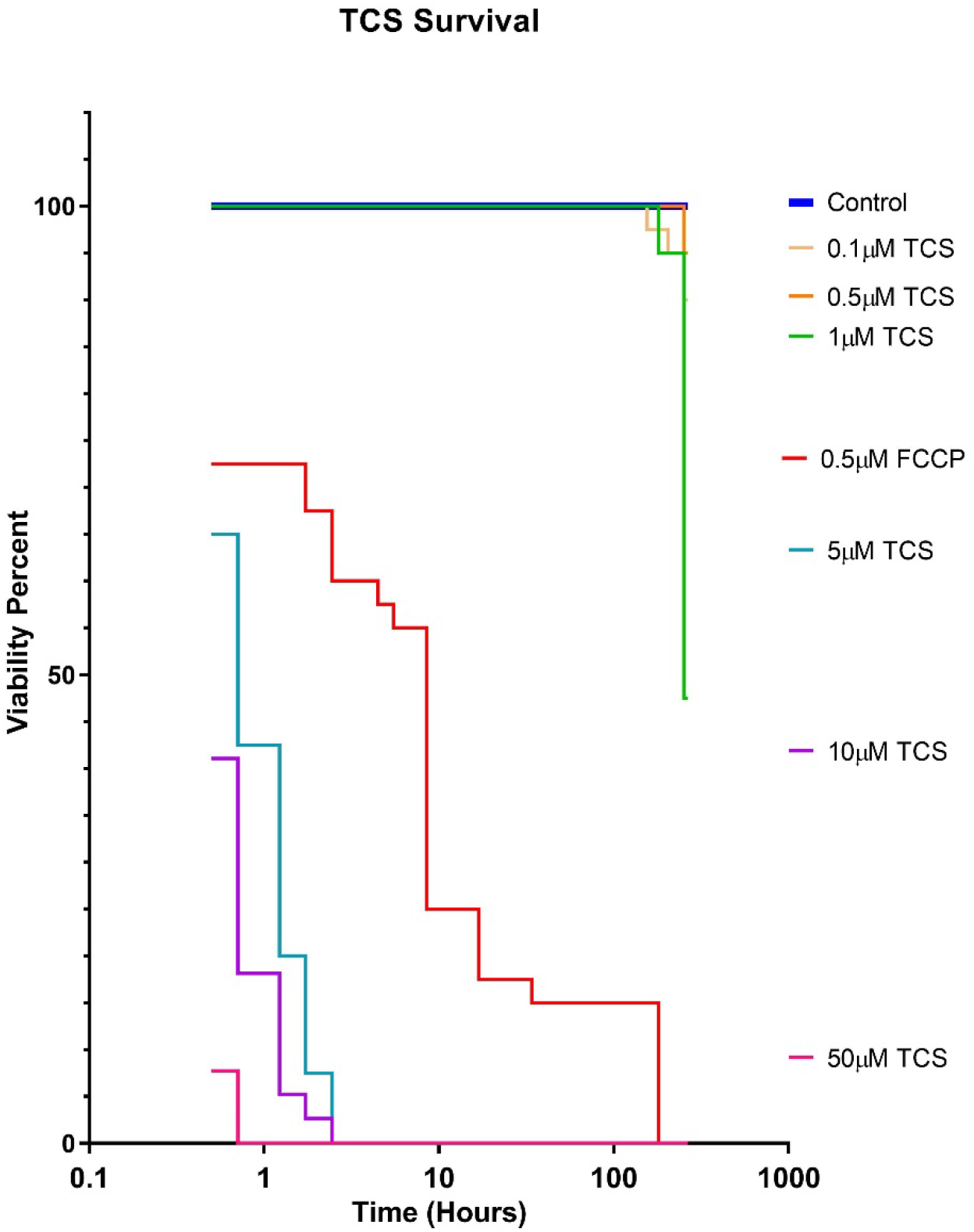
Survival of tadpoles following exposure to TCS and FCCP. Tadpoles exposed to varying concentrations of TCS and FCCP were monitored for survival rates over 14 days. A prolonged exposure time with 1um TCS resulted in significant mortality on the final day of the experiment; however, negligible effects were observed among tadpoles exposed to Control or lower concentrations (0.5 uM and 0.1 uM) of TCS. Concentrations of TCS at 5um and above proved acutely toxic. While FCCP also exhibited toxicity, tadpoles could tolerate a concentration of 0.5 uM for slightly more extended periods.

Confocal microscopy was employed on a Leica SP8 confocal microscope to assess the effects of acute treatment on stage-25 Xenopus laevis tadpoles prior to mortality in the high concentrations used above. Tadpoles were stained with TMRM, a membrane potential dye, to evaluate the mitochondrial activity and membrane potential. For normalization purposes, initial pretreatment images were captured before treating the tadpoles with TCS for 30 minutes while capturing images every 5 minutes. The results of the confocal microscopy analysis showed a significant decrease in mitochondrial activity and membrane potential in tadpoles exposed to TCS compared to the control (see Figure 3 for reference). Where animals treated with 10 uM had roughly a 60% reduction in fluorescence intensity 30 minutes after acute exposure.

**Figure 3:**
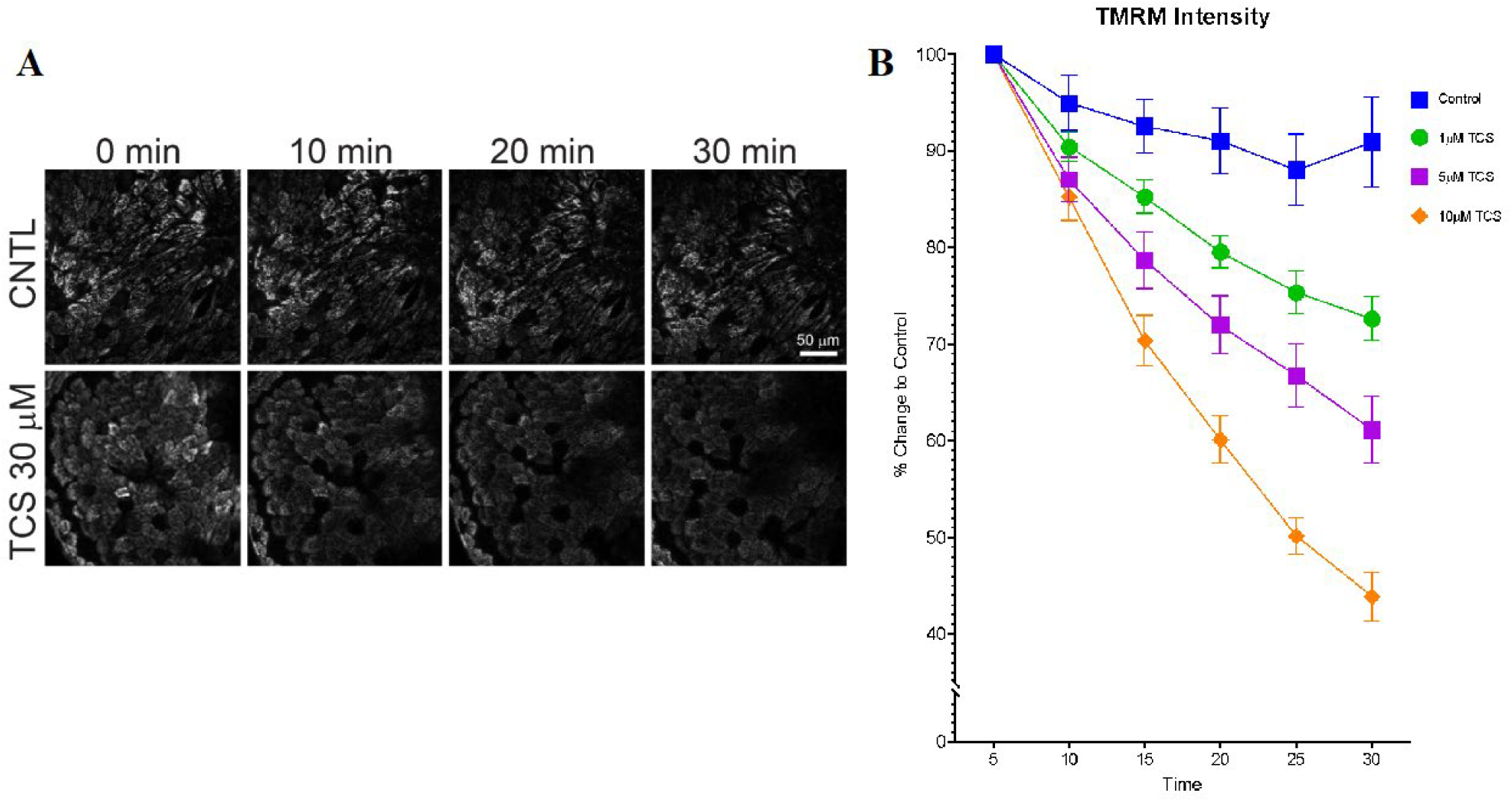
Triclosan (TCS) decreased TMRM intensity in 3-day-old embryos. **A**) Confocal images of TMRM-labeled skin cells in 3-day-old embryos exposed to either control or TCS 30uM show a dramatic decrease in TMRM intensity upon exposure to TCS, indicating a near complete collapse in mitochondrial membrane potential. **B)** Exposure to lower concentrations of TCS (1 uM, 5 uM, and 10 uM) also reduced mitochondrial function, as evidenced by the decreased intensity of TMRM. 10 uM exhibited roughly a 60% decrease in TMRM intensity within 30 minutes of exposure.

Further studies were conducted to investigate the activity of mitochondria in stage-25 Xenopus laevis tadpoles. The XFe24 Seahorse Extracellular Flux Analyzer was used to measure the oxygen consumption rate (OCR) in whole live animals in the XF Islet Plate. Initial OCR measurements were made before the tadpoles were exposed to varying concentrations (30uM, 10uM, or 1uM) of TCS and control solutions. As shown in Figure 4, after exposure to an ATPase inhibitor, oligomycin (10uM), and inner membrane uncoupler FCCP (2.5 uM), OCR levels increased overall for those tadpoles exposed to 30uM TCS even after undergoing oligomycin treatment. This suggests that TCS acts as a mitochondrial uncoupler by inducing loss of ATP-dependent respiration (p <0 .001).

**Figure 4:**
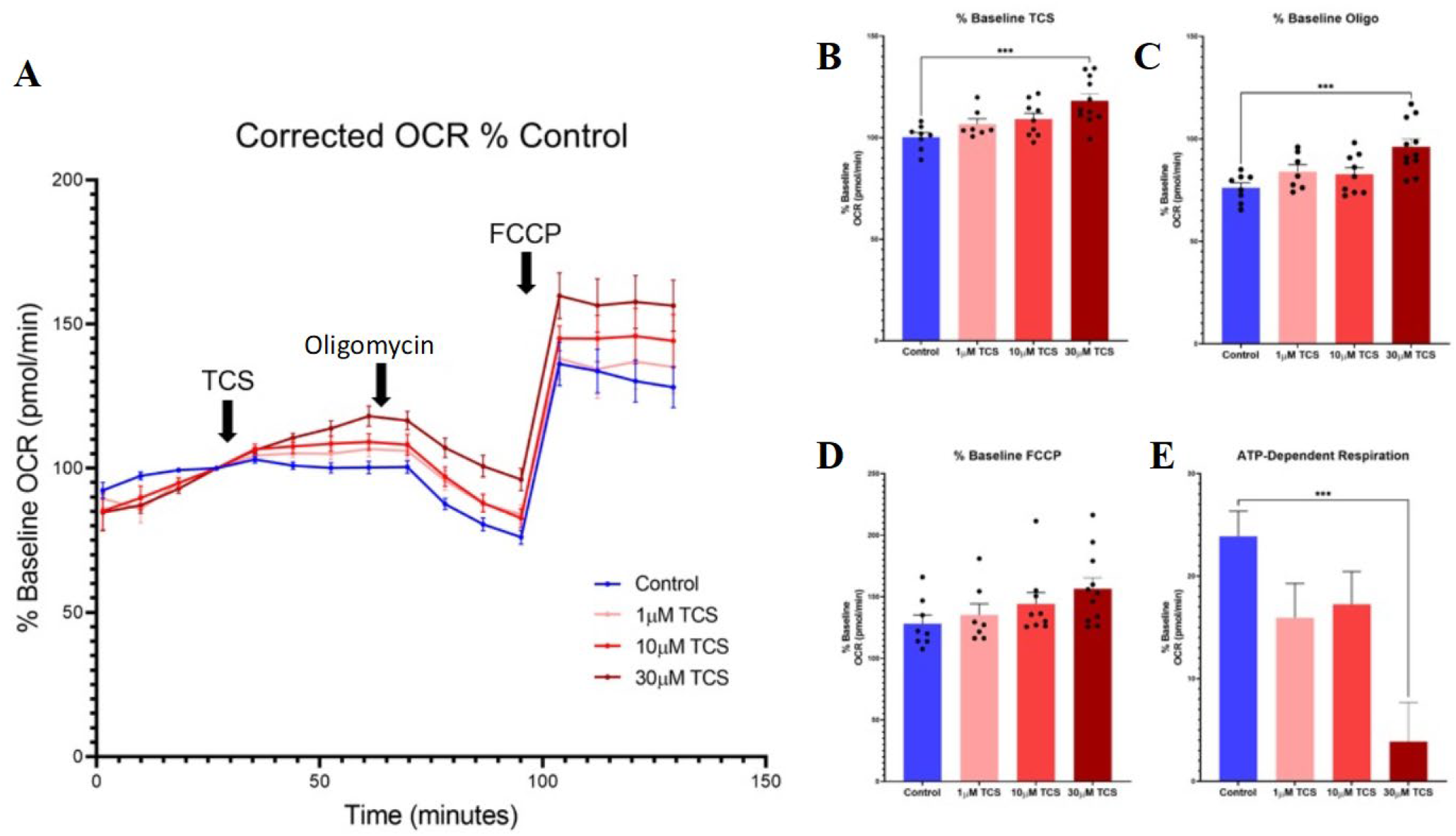
Triclosan increased overall oxygen consumption rate (OCR) but decreased ATP-dependent respiration in whole three-day-old embryos. **A**) OCR was measured in 3-day-old embryos using a Seahorse Extracellular Flux Analyzer. Tadpoles were exposed to control or different concentrations of TCS, followed by ATPase inhibitor (oligomycin) and inner membrane uncoupler (FCCP). **B)** Tadpoles exposed to 30uM TCS significantly increased (***p < 0.001) in OCR following TCS treatment. **C, D)** OCR maintained this significant increase (***p < 0.001) in respiration following the ATPase inhibitor, oligomycin (C), and inner membrane uncoupler, FCCP (D). **E)** When ATP-respiration is calculated by subtracting the normalized oligomycin-induced leak respiration from the TCS basal respiration, a significant decrease (***p < 0.001) in OCR is depicted. Thus, TCS acts as a mitochondrial uncoupler due to the increase in basal and leak (oligomycin) respiration, resulting in a dramatic decrease in ATP-dependent respiration.

To better understand the mechanisms behind TCS-induced mitochondrial dysfunction, we conducted additional confocal imaging experiments using more mature stage-47 tadpoles to examine its impact on developing brains. Intraventricular injections of TMRM were administered to the tadpoles. Following a 24-hour incubation period, tadpoles were exposed to varying concentrations (10uM, 5uM, and 1uM) of TCS or control. The effects were monitored through five-minute intervals via imaging. As shown in Figure 5, our findings revealed a significant reduction in fluorescent intensity observed in TMRM within thirty minutes post-treatment. Additionally, at higher concentrations - particularly those approaching maximum levels used here - induced mitochondrial fission and swelling primarily occurred at endfeet neural progenitor cell sites where differentiation occurs during neurogenesis.

**Figure 5:**
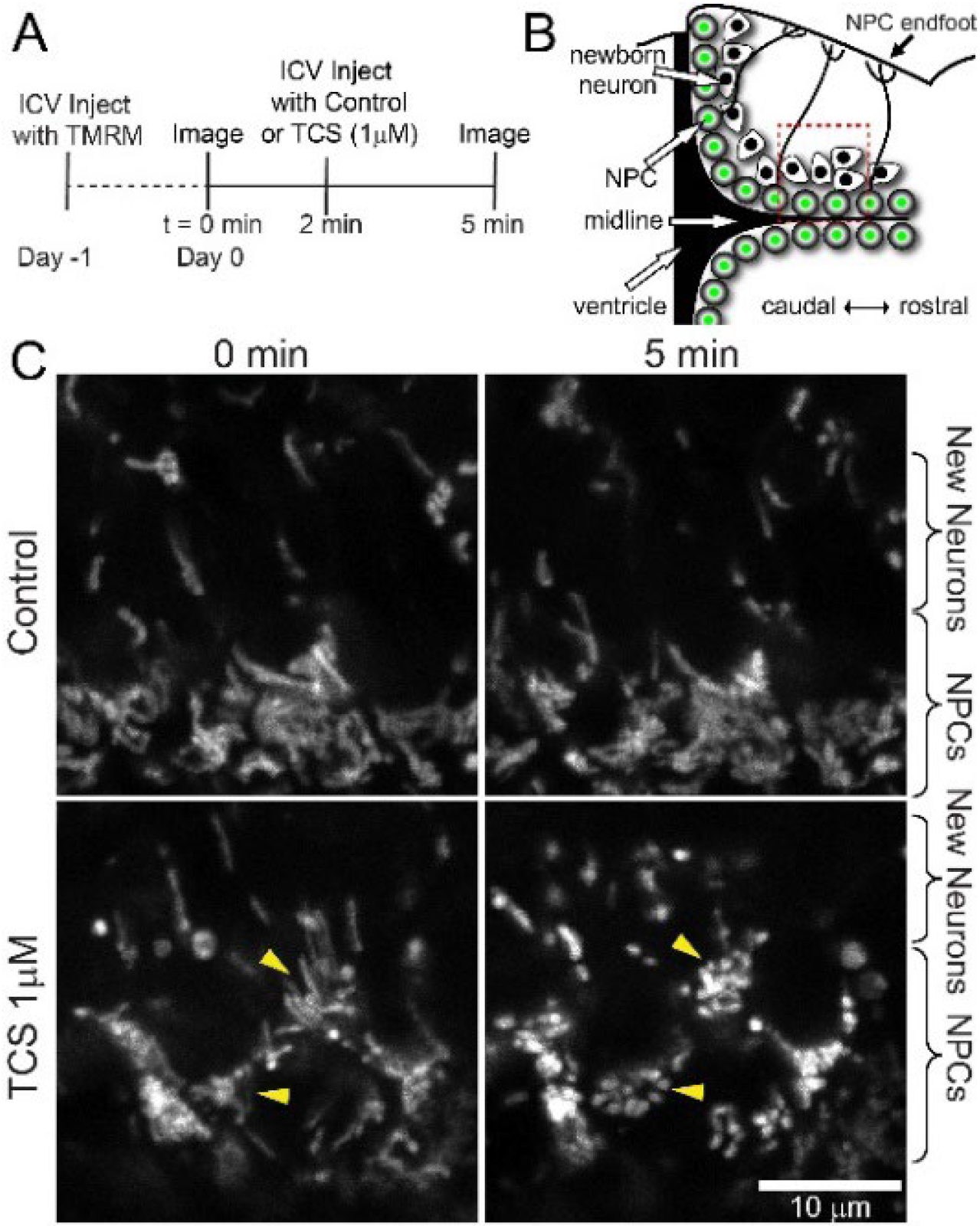
Triclosan decreased TMRM intensity in endfeet of neural progenitor cells and induced mitochondrial fission in Stage 47 tadpoles. We measured TMRM intensity in individual endfeet over time and observed that TCS decreased TMRM intensity. A substantial decrease in TMRM fluorescent intensity is exhibited within 30 min post-treatment. Further, TCS induced mitochondrial fission and swelling at the highest concentrations, particularly in the endfeet of neural progenitor cells. **A)** Tadpoles were injected ICV with TMRM (250 uM). The next day, the ventricular zone was imaged in anesthetized tadpoles. Then the ventricle was injected with either control or 1uM TCS and imaged 3min later. **B**) Illustration showing the imaging location (red box) along the ventricular zone. **C)** Example images showing that TCS induced mitochondrial fission in NPCs (arrowheads) and an apparent increase in TMRM intensity, indicative of an increase in mitochondrial membrane potential.

## Discussion

Initially, TCS was found to cause mitochondrial dysfunction, resulting in adverse cellular consequences such as apoptosis and ATP depletion. As a result, the unregulated and widespread use of TCS can pose an enormous risk to unsuspecting consumers. This risk is further amplified for infants from exposed populations, potentially leading to cognitive deficits and neural stunting. Specifically, we showed that excessive exposure to TCS causes a significant decrease in TMRM fluorescence intensity due to decreased mitochondrial membrane potentials, as observed through confocal microscopy. Further analysis of an XFe24 Seahorse Extracellular Flux Analyzer also showed that OCR levels increased after exposure to 30uM of TCS, indicating that it acts like a mitochondrial uncoupler by inducing loss of ATP-dependent respiration.

Elevated concentrations induced mitochondrial fission and swelling primarily at endfeet neural progenitor cell sites responsible for cell differentiation during neurogenesis within just thirty minutes post-treatment. This is likely an adaptive response to membrane potential loss and the fission pathway’s innate protective mechanism in reducing the spread of pathological counterparts [26]. Fission has been shown to directly cause NPCs to no longer proliferate and differentiate into daughter neurons [27]. This reduces the number of potential daughter neurons and overall synaptic density. These findings suggest that even brief exposure to low concentrations may lead to severe toxicity in developing organisms affecting pivotal physiological processes such as neuronal differentiation via modulating specific aspects of energy metabolism linked with mitochondria activity. Therefore, further research and regulation concerning TCS use are essential as its continued unregulated usage could pose risks beyond developmental implications; this study supports such.

The results obtained from these studies highlight the urgent need for a more comprehensive investigation into the long-term effects of TCS exposure on both human development and environmental systems. The available data indicate that such exposure can significantly affect neurogenesis and cognitive performance, thereby underscoring the necessity of further exploration in this field. Subsequent research endeavors should focus on elucidating the underlying mechanisms of TCS toxicity, with particular emphasis placed upon examining its impact on mitochondrial function and structural alterations, to facilitate the identification of potential therapeutic interventions for individuals exposed to this harmful compound. In light of these potential hazards, global regulations must be implemented regarding TCS usage to prevent manufacturers from endangering future generations through their products.

